# An integrative review of the impact of ungulates on wetland carbon storage and greenhouse gas emissions

**DOI:** 10.1101/2025.10.28.685131

**Authors:** Nadia S. Santini, Edgar J. González, Aymara O. Ramírez-González, Catherine E. Lovelock

## Abstract

Wetlands, spanning freshwater to saline ecosystems, are seasonally or permanently inundated and store over 30% of the world’s carbon within their soils. Ungulates can remove vegetation, disturb soils, and degrade habitats. Although site-level studies exist, broad evaluations of ungulate impacts on carbon stocks and greenhouse gas emissions remain scarce. We assessed how ungulate presence influences aboveground and soil carbon stocks and greenhouse gas emissions in wetlands. Aboveground carbon stocks declined from 36.9 to 3.70 Mg C per ha in the presence of ungulates. Soil carbon stocks were higher in the absence of ungulates (257 Mg C per ha) than in their presence (112 Mg C per ha). CO_2_ equivalent emissions were higher in the presence of non-native ungulates (38.4 Mg CO_2_ eq per ha per yr) than in the presence of native ungulates (27.2 Mg CO_2_ per ha per yr), and higher in freshwater wetlands of warm temperate dry regions when ungulates were present (27.2 Mg CO_2_ eq per ha per yr) compared to when ungulates were absent (13.2 Mg CO_2_ eq per ha per yr). Our findings emphasize the importance of expanding research across different climate regions and soil types to support the development of ungulate management strategies that maintain wetland carbon stocks and reduce greenhouse gas emissions.

## Introduction

Wetlands encompass a wide range of habitats, including marshes, swamps, bogs, fens, ponds, and estuaries (e.g. Kauffman et al. 2004; Kauffman et al. 2016; Zhou et al. 2020). Healthy wetlands support specialized hydrophytic plant communities and provide essential habitats and breeding grounds for animals. Notably, many bird species rely on seasonal wetlands during their migration. In addition to supporting biodiversity, wetlands provide critical ecosystem processes and services, including high levels of vegetative growth and primary productivity, the filtration and supply of water, coastal protection, and flood mitigation (Boughton et al. 2016; Armitage et al. 2019). These critical ecosystem services have a combined estimated global monetary value of US$ 47.4 trillion per year, comprising 44% of the value of services provided by natural ecosystems (Finlayson and Gardner 2021).

Wetlands are covered either seasonally or permanently with fresh, salt, or brackish water, resulting in distinct hydrological processes that include fluctuating water tables, surface water flows, and surface-groundwater interactions (Webb et al. 2012). Due to this prolonged saturation or inundation, wetlands typically have waterlogged or hydric soils with low oxygen levels and high carbon content (Spivak et al. 2019). Accordingly, wetland soils are a critical store of more than a third of the world’s carbon reserves, accumulated through the buildup of detritus over millennia and while they are important global sources of methane (Zhang et al. 2024), they are net greenhouse gas sinks (Iqbal and Shang 2020; Rosentreter et al. 2023). Despite these critical ecological processes and services, global wetland ecosystems face threats from land-use change, urbanization, pollution, invasive species, and sea-level rise, and approximately 35% of wetland area has been lost between 1970 and 2015 (Darra et al. 2019).

A significant threat to wetland ecosystems is the use of wetlands by ungulates such as cattle, buffalo, pigs (and boar), sheep, goats, horses, and donkeys. Most of these species are domesticated and widely introduced beyond their native ranges. They graze vegetation and disturb soils through digging and trampling (pugging), thereby altering wetland vegetation and hydrology (Burdick et al. 2021). Non-native ungulates can also compete with native species for resources, and wild pigs and boar prey on a wide range of species (McDonough et al. 2022). Such impacts make wetlands particularly susceptible to ungulate disturbance, providing the basis for evaluating their effects on carbon storage and greenhouse gas emissions (e.g. Crameri et al. 2025).

The presence of ungulates can result in soil disturbance and erosion, reduced carbon sequestration, and increased greenhouse gas emissions (e.g. Sánchez et al. 2017; Crameri et al. 2025). For example, CO_2_ emissions resulting from soil disturbances caused by wild pigs are suggested to be globally significant (O’Bryan et al. 2022). Nitrous oxide emissions may also rise due to the addition of nitrogen from faeces and urine (Rivera and Chará 2021). The impact of these processes may be influenced by several factors, including the type of ungulate (native or non-native), the intensity of grazing, and the characteristics of the wetland ecosystem, including organic versus mineral soils (Nolte et al. 2015; Davidson et al. 2017).

Ungulate activity can enhance ecosystem degradation, biodiversity loss, and alter nutrient cycling (Morris and Reich 2013; Rowland and Lovelock 2024). For example, wild pigs and boar not only root and damage vegetation but also prey on amphibians, reptiles, invertebrates, and other animal species (e.g. McDonough et al. 2022; Rowland and Lovelock 2024). Conversely, other studies report positive effects, such as enhanced habitat diversity and shifts in plant composition. For instance, in the ephemeral wetlands of California’s Central Valley, sites without grazing animals had 88% more exotic annual grass cover and 44% lower relative cover of native grasses compared to grazed sites (Marty 2005). Similarly, along the French Atlantic coast, grazing was linked to increased plant species richness in wetland communities (Marion et al. 2010). Given the important ecosystem services wetlands provide, and their ongoing degradation, there is a pressing need for wetland restoration, management, and conservation. Measures such as exclusion or adaptive control of domesticated and feral ungulate populations can help reverse wetland loss, maintain carbon sequestration, and preserve the ecological functions that support biodiversity and human well-being (Chausson et al. 2020; Zhu et al. 2020; Lovelock et al. 2022).

In this study, we undertook an integrative review of the impact of ungulates on freshwater and saltwater wetland ecosystems across countries. Our assessment was based on an aggregated analysis of 43 distinct studies spanning from 1987 to 2025. Our objective was to understand the impacts of ungulates on carbon stocks and greenhouse gas emissions in wetland ecosystems. Specifically, we evaluated the influence of ungulates on three key metrics: (i) aboveground carbon stocks, (ii) soil carbon stocks, and (iii) levels of emissions of greenhouse gases (expressed in CO_2_ equivalents, CO_2_ eq), including carbon dioxide (CO_2_), methane (CH_4_), and nitrous oxide (N_2_O). We compared wetland sites with and without ungulates, and evaluated differences in CO_2_ eq across sites with native and non-native ungulates, recognizing that aboveground carbon stocks, soil carbon stocks, and greenhouse gas emissions are influenced by climate zones (Pearson et al. 2015), and soil type (IPCC, 2014; Qin et al. 2014). Our analysis aims to inform management and conservation strategies to enhance the climate change mitigation potential of wetland ecosystems.

## Materials and Methods

We investigated carbon storage and fluxes related to wetland ecosystems’ vegetation, soil, and greenhouse gas emissions. We first compiled published studies that examined the impacts of ungulates, including cattle, sheep, buffalo, goats, pigs, horses, yak, and donkeys, on wetland sites. We selected studies that reported greenhouse gas emissions and carbon stocks, along with aboveground biomass and vegetation cover, that allowed us to determine carbon stocks and emissions. This process resulted in the inclusion of 43 studies from various locations worldwide, covering boreal, temperate, and tropical regions, including studies from the following countries: Australia, Brazil, Canada, China, Denmark, Ecuador, France, India, Mexico, the Netherlands, New Zealand, Peru, Poland, Saudi Arabia, the United Kingdom, and the United States (**Fig. 1a,b**; **Table S1**). Studies included those that used fencing to exclude ungulates (19 studies), compared presence versus absence (22 studies), or managed sites after the exclusion of ungulates (2 studies). These studies often did not report the intensity or activity of ungulates as a quantitative metric, but rather as a qualitative metric (presence and absence, e.g. Hirota et al. 2005; Iram et al. 2021; Lovgren et al. 2022), or when reported, the metric for activity was presented in a variety of forms. In some studies, ungulate activity was quantified as the number of individuals per km^2^ (e.g. Limpert et al. 2021; Xiao et al. 2021; Dalmagro et al. 2022), while in others it was expressed as a rate, such as grazing intensity (Yu and Chmura 2010; Harvey et al. 2019). Because of the variation in approaches used in the literature, we assessed the impacts of the presence versus absence of ungulates across study sites (see **Table S1**).

**Fig 1.**
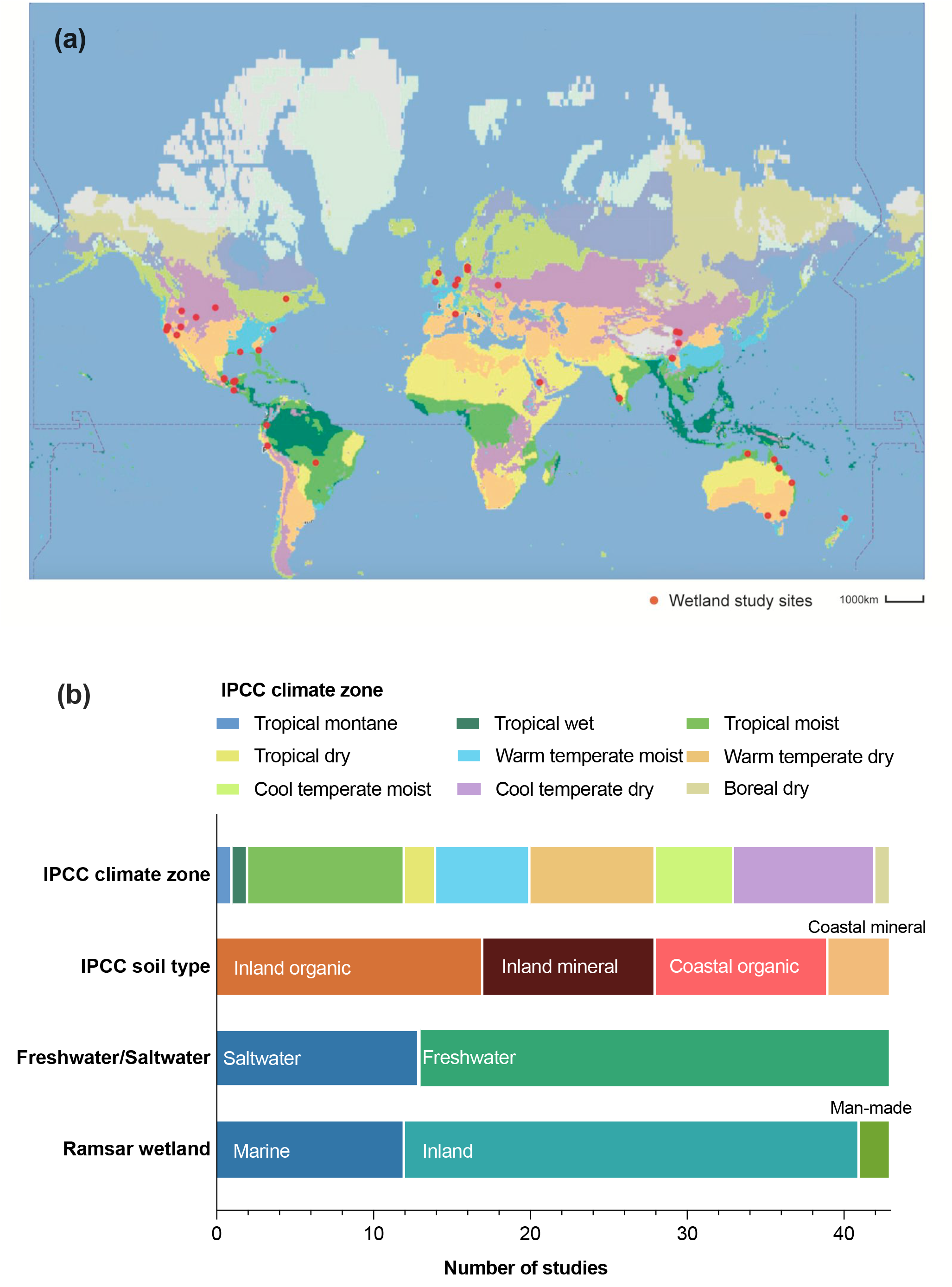
Worldwide study sites used in this integrative review. **(a)** Worldwide map shows the locations of wetland study sites included in this work (precise locations are provided in the Supplementary Materials). **(b)** Studies classified according to climate zone, soil type, and wetland type. Note that each row is independent.

To classify wetland sites, we employed the Ramsar Wetland Classification System, a comprehensive framework designed to recognize the diverse wetland habitats present in each location (Finlayson, 2018). The Ramsar Classification of Wetland Types organizes wetlands into three main groups: marine/coastal wetlands, inland wetlands, and man-made wetlands. This classification system establishes standardized categories for the mapping of wetland sites and ensures consistent terminology in national or regional wetland inventories. In addition, we classified our wetland sites according to the IPCC climate zones (IPCC, 2019). For visualization, we accessed the IPCC climate zones layer from the Google Earth Engine repository (identifier: ‘users/philipaudebert/IPCC/Corrigenda/ipcc_climate_1985-2015_corrigenda_raster’) and applied style parameters to the climate categories. The coordinates of the study sites, derived from ungulate studies, were stored in a featured collection and overlaid on the climate zones (**Fig. 1a, b**). We also classified studies based on wetland soil types according to the 2013 Supplement to the ‘2006 IPCC Guidelines for National Greenhouse Gas Inventories: Wetlands’, as inland mineral soil, coastal mineral soil, inland organic soil, and coastal organic soil (**Fig. 1b; Table S1**; IPCC, 2014). Due to the limited number of studies comprising our dataset, we could not assess the impacts of the presence or absence of ungulates for each climate zone, wetland and/or soil type classification, with the exception of emissions from freshwater wetlands in the warm temperate dry region, where sufficient data allowed for quantitative analysis. A summary of the available data across climate zones, wetland and soil types are presented in **Fig. S1**.

From each study, we either extracted data directly or converted data to obtain aboveground carbon stocks, soil carbon stocks, and greenhouse gas (GHG) emissions. CO_2_ emissions result from the decomposition of organic matter under aerobic conditions. CH_4_ emissions are produced when waterlogged soils create anaerobic conditions, promoting microbial methane production during organic matter decomposition. N_2_O emissions occur because waterlogged soils facilitate denitrification, a microbial process that generates nitrous oxide as a byproduct. For soil carbon stocks, most studies reported data across the first meter of soil. Our analysis included only studies that provided comparable soil carbon stock values based on the presence or absence of ungulates. To account for variability in soil depth, we conducted additional analyses focusing on studies that reported soil carbon stocks specifically for the top 0 – 30 cm layer (see **Fig. S2a–d, Table S2**).

Aboveground biomass data was converted to carbon stocks using a conversion factor of 0.5 and converted to Mg C per ha^-1^ (IPCC, 2006; Myhre et al. 2013). In studies where vegetation cover in percentage was provided (e.g. Elschot et al. 2023), we utilized default estimates for standing biomass based on the IPCC climate zone (IPCC, 2006; IPCC; 2019), where 100% of aboveground biomass cover corresponds to the following values: approximately 1.7 Mg ha^-1^ of dry biomass in the boreal dry and wet zones, and in the cold temperate, dry zone; 2.4 Mg ha^-1^ of dry biomass in the cold temperate, wet zone; 1.6 Mg ha^-1^ of dry biomass in the warm temperate, dry zone; 2.7 Mg ha^-1^ of dry biomass in the warm temperate, wet zone; 2.3 Mg ha^-1^ of dry biomass in the tropical, dry zone, and 6.2 Mg ha^-1^ of dry biomass n the tropical, moist and wet zones.

To enable comparability across studies and sites, we standardized GHG emissions data by converting emission measurements reported in different units to emissions of CO_2_ eq ha^-1^ year^-1^. This standardization was performed in accordance with the IPCC (2014) guidelines. For emissions of CH_4_ and N_2_O, conversions to CO_2_ eq were based on the global warming potential (GWP) values from the IPCC Fifth Assessment Report, where the GWP of CH_4_ is 28, and that of N_2_O is 265 (Myhre et al. 2013; IPCC 2014). GWP quantifies the energy absorbed by the emissions of 1 metric ton (Mg) of a given gas over a specified time period, relative to CO_2_.

When rate of CH_4_ and N_2_O are reported we standardised as:

Mg CH_4_ ha^-1^year ^-1^ x 28 = Mg CO_2_ eq ha^-1^ year^-1^

Mg N_2_O ha^-1^year ^-1^ x 265 = Mg CO_2_ eq ha^-1^ year^-1^

Data standardization enabled to compare aboveground carbon stocks, soil carbon stocks (expressed as Mg C ha^−1^), and greenhouse gas emissions (expressed as Mg CO_2_ eq ha^−1^ year^−1^), across various studies and sites. These comparisons were conducted for both the presence and absence of ungulates across all wetland types (both freshwater and saltwater) and focusing solely on freshwater wetlands. Additionally, we evaluated differences **in** CO_2_ eq **across sites with** native and non-native ungulates and tested for differences in warm temperate dry freshwater wetlands in the presence and absence of ungulates. Given the variability in the provenance of our data, we adopted a conservative approach by employing Hedges’ *g* effect size analysis (Lakens 2013). These comparisons were based on the assessment of mean differences between groups and were designed to correct for potential biases arising from small sample sizes (Hedges 1981; National Institute of Standards and Technology 2017), as: 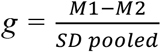; where M_1_ – M_2_ is the difference in means, and SD _pooled_, is the pooled and weighted standard deviation.

Finally, based on statistically significant results (i.e., *p*<0.05), we projected emissions over the next 25 years for all wetlands in the presence of native and non-native ungulates, and for freshwater wetlands in the warm temperate dry region. To estimate the broader impact of ungulate activity on wetland carbon emissions, we projected emissions for a standardized area of 1,000 ha, even though the wetlands studied were often smaller (e.g., ∼100 ha). This approach allows for comparability and scalability of results, facilitating a better understanding of the cumulative effects at landscape and regional scales. To quantify the economic significance of these reductions, we applied a monetary value based on the current carbon market price for CO_2_ eq. For this analysis, we assumed a relatively low carbon price of approximately $20 USD per Mg of CO_2_ eq (World Bank 2023).

## Results

We reviewed a total of 43 studies, including 12 on marine/coastal wetlands, 29 on inland wetlands, and 2 on man-made wetlands. Of these, 17 studies were conducted on inland organic soils, 11 on inland mineral soils, 11 on coastal organic soils, and 4 on coastal mineral soils. Most studies were conducted in tropical moist regions (10), cool temperate dry regions (9), and warm temperate dry regions (8), with no studies reported from boreal moist or polar moist regions (**Fig. 1a,b; Table S1**).

We first evaluated the impact of ungulate presence and absence on aboveground carbon stocks across all wetland sites. In the absence of ungulates, the aboveground carbon stocks were 36.9 ± 15.8 Mg C ha^-1^. Ungulate presence resulted in a large effect size (Hedges’ *g* = 0.70, *p* = 0.006), with aboveground carbon stock being nine times lower, from 36.9±15.8 MgCha^-1^ to 3.70±1.22 MgCha^-1^ across all wetlands (**Fig. 2a, Table S2**). Within freshwater wetlands, aboveground carbon stocks were lower in the presence of ungulates, from 15.9 ± 4.26 Mg C ha^-1^ in the absence of ungulates, to 4.81 ± 1.59 Mg C ha^-1^ in the presence of ungulates (Hedges’ *g* = 0.66, *p*=0.016; **Fig. 2b, Table S2)**.

**Fig 2.**
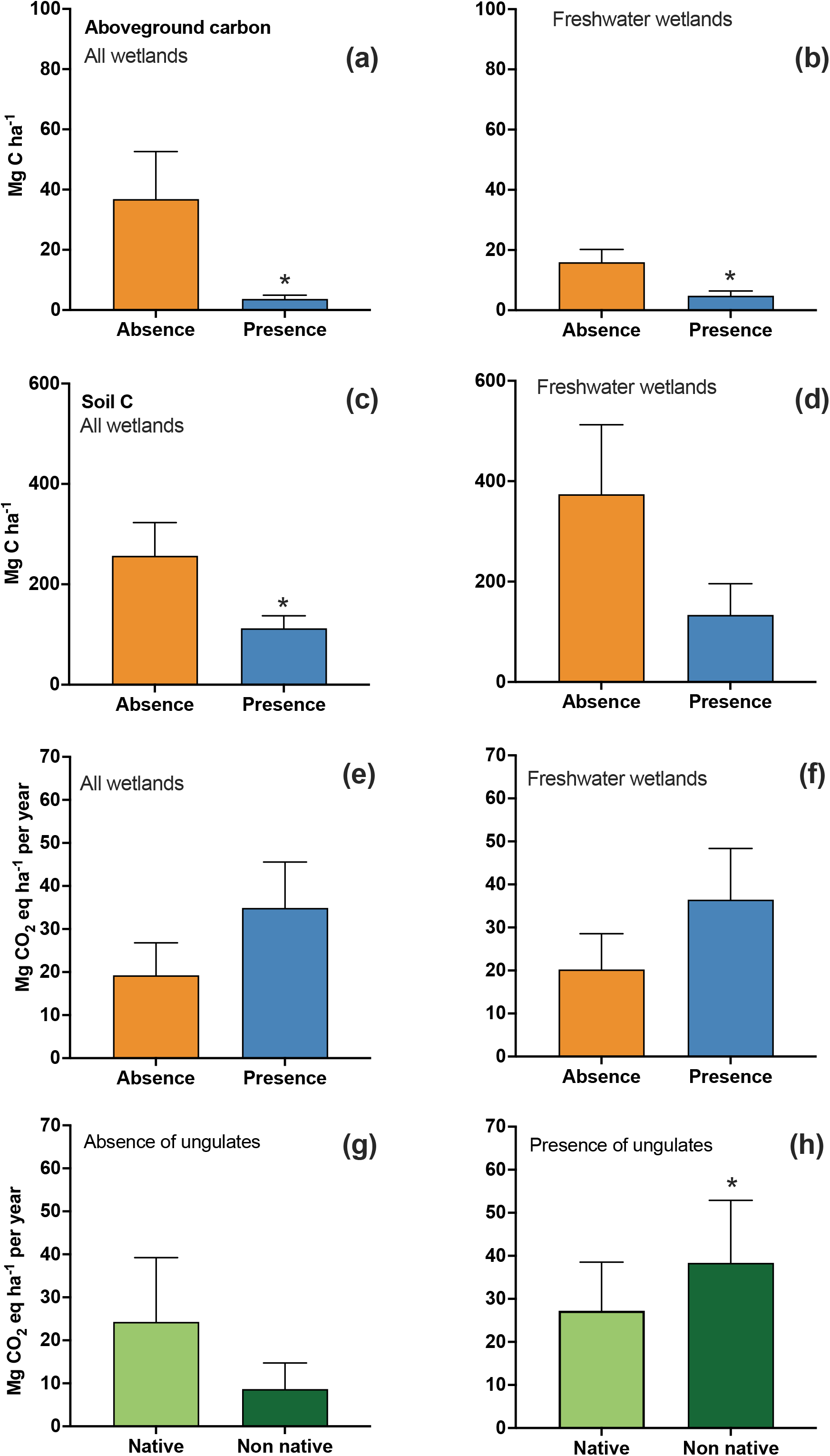
Impact of ungulates on wetlands. **(a)** Aboveground carbon stocks were lower in the presence of ungulates across all wetlands (Hedges’*g* = 0.70, *p*=0.006). **(b)** The presence of ungulates was associated with a significant reduction in the aboveground carbon stocks in freshwater wetlands (Hedges’ *g* = 0.66, *p* = 0.016). **(c)** Soil carbon stocks in the absence and in the presence of ungulates across all wetland sites (Hedges’ *g* = 0.66, *p* = 0.029). **(d)** Soil carbon stocks in the absence and in the presence of ungulates across freshwater wetlands (Hedges’ *g* = 0.75, *p* = 0.052). **(e)** Emissions of CO_2_ eq in the absence and in the presence of ungulates across all wetland sites (Hedges’ *g*, - 0.24, *p*=0.18). **(f)** Emissions of CO_2_ eq in the absence and in the presence of ungulates across freshwater wetland sites (Hedges’ *g*, - 0.24, *p*=0.18). **(g)** Emissions of CO_2_ eq across all wetland sites in the absence of native and non-native ungulates (Hedges’ *g*, 0.32, *p*=0.12). **(h)** Emissions of CO_2_ eq across all wetland sites in the presence of native and non-native ungulates (Hedges’ *g* = - 0.15, *p*=0.038).

We compared soil carbon stocks in wetlands and observed that, in the presence of ungulates, soil carbon stocks were 112 ± 24.9 Mg C ha^−1^ compared to 257 ± 66 Mg C ha^−1^ in the absence of ungulates across all wetlands (Hedges’ *g* = 0.66, *p*=0.029), and 133.5 ± 62.3 Mg C ha^−1^ compared to 374 ± 138 Mg C ha^−1^ in the absence of ungulates in freshwater wetlands (Hedges’ *g* = 0.75, *p*=0.052, **Fig. 2c,d**; **Table S2**).

We compared the CO_2_ eq emissions across all wetland sites in the presence or absence of ungulates, and found that when ungulates were absent, global wetlands emit an average of 19.2 ± 7.54 Mg CO_2_ eq ha^-1^ year^-1^, whereas wetlands in the presence of ungulates emit 34.9 ± 10.7 Mg CO_2_ eq ha^-1^ year^-1^. The effect size Hedges’ *g* was calculated to be - 0.24 (*p*=0.18; **Fig. 2e, Table S2**). For freshwater wetlands, emissions associated with the presence of ungulates were 36.5 ± 11.9 Mg CO_2_ eq ha^-1^ year^-1^, compared to average emissions of 20.2 ± 8.31 Mg CO_2_ eq ha^-1^ year^-1^ in the absence of ungulates (Hedges’ *g* = -0.24, *p*=0.18, **Fig. 2f, Table S2**). We also compared CO_2_ eq emissions across sites where native and non-native ungulates were absent and present. Within sites where both native and non-native ungulates were absent, emissions tended to be higher in sites where native ungulates naturally occur (24.3 ± 14.9 Mg CO_2_ eq ha^-1^ year^-1^), compared to sites where non-native ungulates were present (8.65 ± 6.08 Mg CO_2_ eq ha^-1^ year^-1^, Hedges’*g =* 0.32, *p*=0.12, **Fig. 2g, Table S2**). In the presence of ungulates, emissions were significantly larger when non-native ungulates were present (38.4 ± 14.5 Mg CO_2_ eq ha^-1^ year^-1^), than when native ungulates were present (27.2 ± 11.3 Mg CO_2_ eq ha^-1^ year^-1^, Hedges’*g =* - 0.15, *p*=0.038, **Fig. 2h, Table S2**).

Overall, we found that CO_2_ eq emissions from wetlands varied among climate zones (**Fig. S1a, b**). The limited data for each climate zone and soil type limited our ability to conduct further quantitative analyses for most categories. However, for emissions from freshwater wetlands in the warm temperate dry region, sufficient data were available, enabling us to perform a detailed quantitative analysis. We found that in the presence of ungulates across this region, emissions were significantly larger (27.2 ± 4.05 Mg CO_2_ eq ha^-1^ year^-1^) than when ungulates were absent (13.2 ± 2.86 Mg CO_2_ eq ha^-1^ year^-1^; Hedges’*g =* -1.42, *p*=0.04, **Table S2**).

We observed that mean CO_2_ eq emissions from wetlands were highest in organic soils, compared to mineral soil types, however, due to high variability in the data and limited sample sizes, further statistical analyses were not performed (**Fig. S1c**).

Finally, by comparing the projected cumulative CO_2_ eq emissions over 25 years from 1,000 hectares of wetlands in the presence of native and non-native ungulates, we found there was a 29% reduction in CO2 eq emissions in the presence of native versus non-native ungulates, totalling 280,000 Mg CO2 eq (**Fig. 3a)**. Assuming a relatively low carbon price ∼$20 USD per Mg CO_2_ eq, removing non-native ungulates from 1,000 ha of wetlands for 25 years would have an estimated value of $5,600,000 USD. For freshwater wetlands across the warm temperate dry climatic zone, cumulative CO_2_ eq emissions were 330,000 Mg CO_2_ eq at sites where ungulates are absent, and 680,000 Mg CO_2_ eq in the presence of ungulates (**Fig. 3b)**.

**Fig 3.**
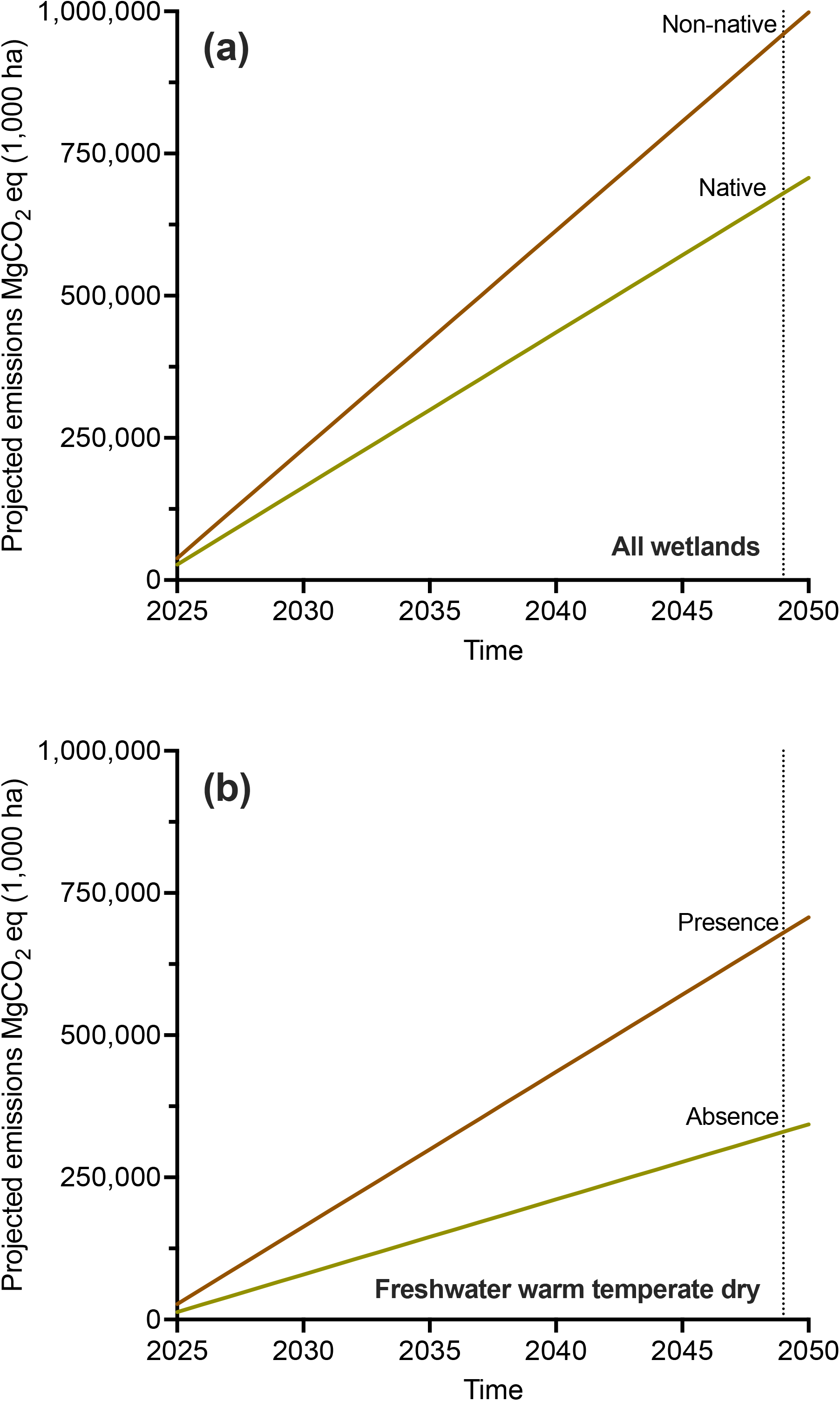
Projected impact of ungulates on carbon emissions over 25 years. **(a)** Cumulative CO_2_ eq emissions over 25 years across 1,000 ha in the presence of native and non-native ungulates were 680,000 Mg CO_2_ eq, and 960,000 Mg CO_2_ eq, respectively. **(b)** Across warm temperate dry wetlands, cumulative CO_2_ eq emissions over 25 years across 1,000 ha in the absence of ungulates was 330,000 Mg CO_2_ eq, and 680,000 Mg CO_2_ eq in the presence of ungulates.

## Discussion

Our analysis revealed a geographic bias in the available studies of the impacts of ungulates on wetland carbon cycling. Most of the studies (29) focused on inland freshwater wetlands, while only 12 assessed the impact of ungulates on carbon cycling in marine/coastal wetlands, and 2 on man-made wetlands (Lovgren et al. 2022; Elschot et al. 2023). The underrepresentation of marine wetlands could be due to the more limited documented use of these wetlands for grazing outside Europe, where livestock grazing in saltmarshes is common (e.g. Davidson et al. 2017) and due to limited sampling of the influence of ungulates on saline wetlands in other regions (e.g. Prahalad et al. 2024). Grazing in man-made wetlands was only studied in Europe and the United States, highlighting the need for further research to better estimate the impacts of ungulate presence on carbon cycling in these environments. We identified a significant research gap in boreal and polar wetlands. These ecosystems are highly sensitive to climate change and are expected to experience accelerated warming, making further research important for understanding the impact of native and non-native ungulates in the global carbon cycle.

Our analysis confirmed that the presence of ungulates significantly reduces aboveground carbon stocks in wetlands. Grazing by ungulates directly consumes plant biomass, leading to reductions in carbon stored in vegetation (Doupé et al. 2010; Limpert et al. 2021) with potential to have flow-on effects on carbon storage in soils (e.g. Zhang et al. 2002). Declines in vegetative cover can also cause long-term shifts in plant community composition, often resulting in the dominance of stress-tolerant, less palatable species (Jones et al. 2011; Rowland and Lovelock 2024). The species composition of a wetland community is important for ecosystem resilience, as the complementary use of resources and functional traits of different species can enhance wetland productivity and the ability to recover from disturbance (Liu et al. 2022). However, shifts in vegetation composition following disturbance may persist for years or even decades, potentially reducing carbon sequestration, wetland productivity, and resilience to stressors such as climate change and land-use changes (Yang et al. 2022).

We found that sites with non-native ungulates were associated with significantly higher CO_2_ eq emissions compared to sites with native ungulates. In wetlands, soil disturbance from non-native species, can enhance the release of stored carbon through enhanced soil aeration and organic matter decomposition (Lovgren et al. 2022; Elschot et al. 2023). This difference may be attributed to several factors. For example, non-native species often have higher stocking densities, which can lead to greater soil disturbance, compaction, or erosion (O’Bryan et al. 2022). Additionally, non-native ungulates frequently alter plant community composition, which can reduce the wetlands’ capacity to sequester carbon in some sites (Rowland and Lovelock 2024, and references therein). In contrast, native ungulates, have co-evolved within their ecosystems, and tend to exert less disruptive grazing pressure (e.g. Allen et al. 2023). Additionally, we found that ungulate presence was associated with significantly higher CO_2_-eq emissions in freshwater wetlands of the warm temperate dry climate zone. In these sites, soil disturbance caused by trampling, rooting, or grazing can enhance the release of stored carbon through increased aeration and organic matter decomposition (Lovgren et al. 2022; Elschot et al. 2023). This effect was only detected in warm temperate dry wetlands, likely due to the limited number of studies available for other climate zones. With more studies across regions and wetland types, it may possible that similar or even stronger differences in greenhouse gas emissions would emerge.

Because of the limited availability of data, we could not assess how contributions of different greenhouse gases to overall emissions varied among different climate zones, wetland and soil types. Our findings highlight the need for further studies that systematically compare the contributions of CH_4_ and N_2_O over a range of sites (encompassing different climate zones, wetland, and soil types) with and without ungulates to better quantify the global impact of native and non-native ungulates on greenhouse gas emissions from wetlands.

Our analysis did not allow for testing of the effects of ungulate density on carbon stocks or emissions, or allow for specific recommendations regarding management of ungulate densities to enhance carbon stocks and reduce emissions. This is because most studies included in our review reported ungulate impacts as presence or absence rather than in response to a known density of animals (e.g., Hirota et al. 2005; Iram et al. 2021; Lovgren et al. 2022). We recommend that future studies on the impacts of ungulates on wetland carbon cycling standardize measures of ungulate density to enable analysis that could support development of ungulate management strategies and models that could be focussed on thresholds of animal densities (e.g. Carpio et al. 2021; O’Bryan et al. 2022).

Our data suggested that the type of soils may play a role in determining the magnitude of greenhouse gas emissions due to ungulate presence. Organic soils, which have higher organic carbon content compared to mineral soils, have been observed to have greater emissions when disturbed compared to mineral soil types (Lovelock et al. 2017). Ungulates can disturb carbon-rich soils as they forage, accelerating microbial decomposition of organic matter and releasing greenhouse gases into the atmosphere. In contrast, mineral soils, though still affected by ungulate activity, likely exhibit comparatively lower emissions due to their lower organic matter content, although direct tests of this hypothesis are still needed.

Finally, by comparing the projected cumulative CO_2_ eq emissions over 25 years from 1,000 hectares of wetlands, we found a 29% reduction in CO_2_ eq emissions in the presence of native versus non-native ungulates, totalling 280,000 Mg CO2 eq (i.e., an annualized per hectare CO_2_ emission of 9 Mg CO_2_ eq ha^-1^ year^-1^). O’Bryan et al. (2022) estimated global CO_2_ emissions due to wild pig soil disturbance to range from 0.40 to 3.19 Mg CO_2_ eq ha^−1^ year^−1^. However, their analysis did not specifically focus on wetlands but rather encompassed multiple land types. Additionally, their estimates accounted only for CO_2_ emissions, excluding potential contributions from methane (CH_4_) and nitrous oxide (N_2_O). Future research should further refine local and global models to better quantify the full scope of emissions associated with ungulate impacts.

Assuming a relatively low carbon price of ∼$20 USD per Mg CO_2_ eq, the reductions in emissions from managing non-native ungulates would generate an estimated value of $5,600,000 USD over 25 years. Thus, the benefits of managing non-native ungulate impacts on wetlands could be sufficient to develop carbon projects with economic benefits that could provide funding for conservation and restoration of wetlands (e.g. Lovelock et al. 2022). The VM0033 Methodology for tidal wetland and seagrass restoration (VERRA, 2023) allows activities that include “Improving management practice(s) (e.g., removing invasive species, reduced grazing)”. Additionally, the Gold Standard Blue Carbon and Freshwater Wetlands methodology (Gold Standard, 2024) includes “Natural regeneration of native vegetation by improving management practices (e.g., removing invasive species, improving tidal channels)” for coastal wetlands, and for freshwater wetlands. Ungulate management may therefore be accommodated within these carbon market methodologies.

## Conclusions

While our study provides valuable insights into the effects of ungulates on wetland carbon dynamics, further research is needed to address key knowledge gaps. There is a need for more studies in underrepresented regions such as wetlands in boreal, polar regions, and for saline wetlands, as well as man-made wetland systems. Additionally, more systematic research is required to evaluate the effectiveness of different management strategies, such as rotational grazing and exclusion of ungulates on carbon stocks and greenhouse gas emissions from wetlands. Research that establishes linkages between ungulate density and greenhouse gas emissions are also needed. Our integrative review, suggests that ungulate management may serve as a potential component of carbon projects aimed at mitigating emissions through wetland conservation or restoration. Regional variation in emissions due to ungulate type, climate and wetland soil type should be considered when evaluating levels of emissions and the financial feasibility of carbon projects implementing ungulate management in wetland ecosystems.

## Data availability

Data underlying this study, compiled from previously published sources, are available from the corresponding author upon reasonable request. A summary of the compiled datasets is provided in the Supplementary Materials.

