## Supplementary Materials for "An integrative review of the impact of ungulates on wetland carbon storage and greenhouse gas emissions"

| **Table S1.** Summary of studies used in this comparative analysis. SOC: Soil Organic Carbon. The global warming potential (GWP) for CH_4_ was taken as 28, while the GWP for N_2_O was determined to be 265, climate zone and soil type were classified according to the IPCC guidelines (IPCC, 2006; 2014; Myhre *et al.,* 2013). Ungulate activity is reported as in the original manuscripts. N/A not available. | | | | | | | | | | | |
| --- | --- | --- | --- | --- | --- | --- | --- | --- | --- | --- | --- |
| **Country** | **Continent** | **Ramsar wetland type** | | **IPCC climate zone,**  **soil type** | **Temperature**  **range (°C)** | **Rainfall**  **(mm)** | **Type of study** | **Ungulate type (native or non-native)** | **Ungulate**  **activity** | **Data recorded** | **Reference** |
| Australia | Oceania | Inland wetland | Freshwater | Warm temperate dry, inland mineral | 14 to 28 | 500 | Exclusion fencing | Sheep (non-native) | 9.4 sheep ha^-1^ | CO_2_  Aboveground C stock | Limpert *et al.,* (2021) |
| Australia | Oceania | Inland wetland | Freshwater | Tropical wet, inland mineral | 14 to 23 | 2,158 | Presence and absence of ungulates | Cattle **(**non-native) | N/A | CO_2_  CH_4_ | Iram *et al.,* (2021) |
| Australia | Oceania | Inland wetland | Freshwater | Tropical moist, inland organic | 18 to 27 | 1,200 | Exclusion fencing | Pigs **(**non-native) | N/A | Vegetation cover | Doupé *et al*., (2010) |
| Australia | Oceania | Inland wetland | Freshwater | Warm temperate dry, inland organic | 2.8 to 32 | 544 | Managed after exclusion | Sheep **(**non-native) | N/A | Aboveground biomass | Treby *et al.,* (2020) |
| Australia | Oceania | Inland wetland | Freshwater | Tropical moist, inland organic | 17 to 34 | 1,400 | Exclusion fencing | Buffalo, pigs **(**non-native) | N/A | Vegetation cover | Sloane *et al.,* (2021) |
| Australia | Oceania | Inland wetland | Freshwater | Cool temperate dry, inland organic | -3.5 to 22 | 1,816 | Presence and absence of ungulates | Horses **(**non-native) | N/A | CO_2_ | Treby and Grover (2023) |
| Australia | Oceania | Inland wetland | Freshwater | Tropical moist, inland organic | 17 to 34 | 1,400 | Presence and absence of ungulates | Buffalo **(**non-native) | When absent:  vegetation biomass 5 to 8 Mg ha ^-1^  When present:  2 to 3 Mg ha^-1^ | Vegetation cover | Skeat *et al.*, (1996) |
| Australia | Oceania | Marine / Coastal wetland | Saltwater | Tropical dry, coastal organic | 7 to 33 | 1,306 | Exclusion fencing | Pigs, cattle **(**non-native) | N/A | SOC | Waltham , *et al.* (2023) |
| Australia | Oceania | Marine / Coastal wetland | Saltwater | Tropical moist,  coastal organic | 23 to 31 | 1,600 | Exclusion fencing | Pigs and buffalo (non-native) | N/A | CO_2_  CH_4_  SOC | Crameri *et al*. (2025) |
| Brazil | America | Inland wetland | Freshwater | Tropical moist, inland mineral | 21 to 29 | 1,345 | Presence and absence of ungulates | Cattle  **(**non-native) | Variable, 0.29 to 0.69 head ha^-1^. | CO_2_ | Dalmagro *et al.,* (2022) |
| Canada | America | Marine / Coastal wetland | Saltwater | Cool temperate moist, coastal organic | -13 to 23 | 1,469 | Presence and absence of ungulates | Sheep  **(**non-native) | 100 lambs grazed for at least six hours per day for 90 days. | Aboveground biomass | Yu, Chmura (2010) |
| China | Asia | Inland wetland | Freshwater | Cool temperate dry, inland organic | -10 to 10 | 650 | Presence and absence of ungulates | Endemic fossorial vertebrates (native) | N/A | CO_2_  CH_4_  N_2_O | Zhou *et al.,* (2020) |
| China | Asia | Inland wetland | Freshwater | Boreal dry, inland mineral | -6 to 9 | 500 | Exclusion fencing | Sheep, yak, cattle (native, non-native) | N/A | CO_2_  CH_4_ | Hirota *et al*., (2005) |
| China | Asia | Inland wetland | Freshwater | Tropical moist, inland mineral | -4 to 13 | 620 | Presence and absence of ungulates | Tibetan pig (native) | 16 yaks km^-2^  60 sheep km^-2^ | SOC | Xiao *et al*., (2021) |
| China | Asia | Inland wetland | Freshwater | Cool temperate dry, inland mineral | -4 to 13 | 620 | Presence and absence of ungulates | Tibetan pig, yak  (native) | 115 km^-2^ | Aboveground biomass | Wang *et al*., (2018) |
| China | Asia | Inland wetland | Freshwater | Cool temperate dry, inland organic | -2.2 to 24 | 610 | Exclusion fencing | Sheep, yak  (native, non-native) | 2.14 sheep equivalents ha^-1^ | CH_4_ | Dang *et al*., (2023) |
| Denmark | Europe | Marine / Coastal wetland | Saltwater | Cool temperate moist, coastal organic | 7 to 11 | 651 | Presence and absence of ungulates | Cattle  (native) | N/A | Aboveground biomass  SOC | Lovgren *et al.,* (2022) |
| Ecuador | America | Inland wetland | Freshwater | Warm temperate moist, inland organic | 7 to 18 | 1,600 | Presence and absence of ungulates | Cattle  (non-native) | N/A | CO_2_  CH_4_ | Sánchez *et al*., (2017) |
| France | Europe | Marine / Coastal wetland | Saltwater | Warm temperate dry, coastal mineral | 0 to 28 | 625 | Presence and absence of ungulates | Cattle  (native) | High: presence of cattle during summertime when the capacity to alter carbon dynamics peaks.  Medium:  Presence of cattle during winter, with less capacity to alter carbon dynamics).  Absent: current absence. | C sequestration rate | Martínez-Eixarch *et al.* (2024) |
| India | Asia | Inland wetland | Freshwater | Tropical montane, inland organic | 0 to 20 | 1,887 | Presence and absence of ungulates | Buffalo, cattle  (native) | Considered as a continuous descriptive variable. | Vegetation cover | Mohandass *et al.,* (2016) |
| Mexico | America | Marine / Coastal wetland | Saltwater | Tropical moist, coastal organic | 16 to 20 | 1,693 | Presence and absence of ungulates | Cattle  (non-native) | N/A | Aboveground biomass  SOC | Kauffman *et al.,* (2016) |
| Mexico | America | Inland wetland | Freshwater | Tropical moist, inland organic | 13 to 32 | 1,425 | Presence and absence of ungulates | Cattle  (non-native) | 2 to 3 head ha^-1^ | Aboveground carbon stock  SOC | Hernández *et al*., (2015) |
| Mexico | America | Inland wetland | Freshwater | Tropical moist, inland organic | 18 to 36 | 1,425 to 1,750 | Presence and absence of ungulates | Cattle  (non-native) | N/A | Aboveground carbon stock | Sjogersten *et al*., (2021) |
| Netherlands | Europe | Man-made wetland | Freshwater | Warm temperate moist, inland organic | 8 to 18 | 793 | Presence and absence of ungulates | Cattle  (native) | Presence described as intensive rotational grazing | CO_2_ | Veenendaal *et al.,* (2007) |
| Netherlands | Europe | Marine / Coastal wetland | Saltwater | Warm temperate moist, coastal organic | 5 to 23 | 780 | Exclusion fencing | Cattle  (native) | 20 to 25 head km^-2^ | Vegetation cover | Elschot *et al*., (2023) |
| New Zealand | Oceania | Inland wetland | Freshwater | Warm temperate moist, inland organic | 9 to 19 | 1,199 | Exclusion fencing | Cattle  (non-native) | N/A | CO_2_ | Campbell *et al.,* (2015) |
| Peru | America | Inland wetland | Freshwater | Cool temperate dry, inland organic | 0 to 24 | 800 | Presence and absence of ungulates | Cattle  (non-native) | N/A | SOC | Chimner *et al.,* (2023) |
| Poland | Europe | Inland wetland | Freshwater | Cool temperate dry, inland organic | 0 to 30 | 575 | Exclusion fencing | Horses  (native) | 0.1 large stock unit ha^-1^ | Vegetation cover | Chodkiewicz *et al.,* (2023) |
| Saudi Arabia | Asia | Marine / Coastal wetland | Saltwater | Tropical dry, coastal organic | 30 to 34 | 81 | Presence and absence of ungulates | Camels  (native) | N/A | SOC  C sequestration rate | Dajam *et al.,* (2024) |
| United Kingdom | Europe | Inland wetland | Freshwater | Cool temperate moist, inland organic | -1.5 to 24.5 | 1,900 | Exclusion fencing | Sheep  (non-native) | N/A | Aboveground Carbon Stock  SOC | Ward *et al*., (2007) |
| United Kingdom | Europe | Marine / Coastal wetland | Saltwater | Cool temperate moist, coastal organic | 5 to 25 | 1,382 | Presence and absence of ungulates | Sheep, cattle  (native) | 0.3 – 0.7 large stock unit ha^-1^  year^-1^ | Aboveground biomass | Harvey *et al.*, (2019) |
| United States of America | America | Inland wetland | Freshwater | Warm temperate dry, inland mineral | 10 to 27 | 708 | Exclusion fencing | Cattle  (non-native) | Presence as 50% percentage of biomass removed | CH_4_ | Oates *et al*., (2008) |
| United States of America | America | Man-made wetland | Saltwater | Warm temperate dry, coastal mineral | 7 to 30 | 325 | Presence and absence of ungulates | Cattle  (non-native) | Day and night presence of cattle | CH_4_ | Baldocchi *et al*., (2012) |
| United States of America | America | Inland wetland | Freshwater | Cool temperate dry, inland mineral | 1 to 10 | 308 | Managed after exclusion | Cattle  (non-native) | 0.8 animal unit per month | CO_2_  CH_4_ | Finocchiaro *et al.,* (2014) |
| United States of America | America | Inland wetland | Freshwater | Tropical moist, inland organic | -5 to 18 | 1,200 | Exclusion fencing | Cattle  (non-native) | N/A | Aboveground Carbon Stock | Kauffman *et al.,* (2004) |
| United States of America | America | Marine / Coastal wetland | Saltwater | Warm temperate moist, coastal organic | 15 to 32 | 1,600 | Exclusion fencing | Nutria and wild boar  (non-native) | Nutria values reported as 22 ha^-1^, NA for wild boar | Aboveground biomass | Ford and Grace (1998) |
| United States of America | America | Inland wetland | Freshwater | Warm temperate dry, inland mineral | 8 to 14 | 420 | Exclusion fencing | Cattle  (non-native) | Animal biomass of 1,000 kg ha^-1^ - 1,500 kg ha^-1^ | CH_4_ | Allen-Diaz *et al.,* (2004) |
| United States of America | America | Inland wetland | Freshwater | Warm temperate dry, inland mineral | 8 to 14 | 420 | Exclusion fencing | Cattle  (non-native) | Animal biomass of 1,000 kg ha^-1^ - 1,500kg ha^-1^ | Aboveground biomass | Allen-Diaz and Jackson (2000) |
| United States of America | America | Inland wetland | Freshwater | Cool temperate dry, inland organic | -1 to 10 | 330 | Exclusion fencing | Sheep, cattle  (non-native) | Presence described as activity for more than 20 days annually | Aboveground biomass | Booth *et al*., (2022) |
| United States of America | America | Inland wetland | Freshwater | Warm temperate dry, inland mineral | -5 to 55 | 50 | Presence and absence of ungulates | Donkey  (non-native) | N/A | Vegetation cover | Lundgren *et al.,* (2022) |
| United States of America | America | Marine / Coastal wetland | Saltwater | Warm temperate moist, coastal mineral | 0.6 to 6.8 | 406 | Exclusion fencing | Horses  (non-native) | N/A | Vegetation cover | Furbish and Albano (1994) |
| United States of America | America | Marine / Coastal wetland | Saltwater | Warm temperate moist, coastal mineral | 12 to 28 | 1,340 | Exclusion fencing | Horses  (non-native) | N/A | Aboveground biomass | Turner (1987) |
| United States of America | America | Inland wetland | Freshwater | Cool temperate dry, inland organic | -10 to 30 | 300 | Presence and absence of ungulates | Horses, sheep, cattle (non-native) | Continuous from 0 – 6 animal unit per month | Vegetation cover | Burdick *et al.*, (2021) |

**Table S2**. Mean, standard error (SE), sample size (n _mean_) that refers to the number of values used in our integrative review, Hedges' *g*, and *p*-values for the studied parameters across the included studies.

| **Assessed parameter** | **Absence** | | | **Presence** | | | **Hedges’*g*** | ***p*-value** |
| --- | --- | --- | --- | --- | --- | --- | --- | --- |
|  | **mean** | **SE** | **n _mean_** | **mean** | **SE** | **n _mean_** |  |  |
| Aboveground C stock (all wetlands) | 36.9 | 15.8 | 15 | 3.7 | 1.22 | 12 | 0.70 | 0.006 |
| Aboveground C (freshwater) | 15.9 | 4.26 | 28 | 4.81 | 1.59 | 33 | 0.66 | 0.016 |
| Soil C stock (all wetlands, 0 – 30 cm and 0 – 100 cm) | 257 | 66.0 | 19 | 112 | 24.9 | 22 | 0.66 | 0.029 |
| Soil C stock (freshwater, 0 – 30 cm and 0 – 100 cm) | 374 | 138 | 8 | 133 | 62.3 | 8 | 0.75 | 0.052 |
| Soil C stock (0 – 30 cm, all wetlands) * | 104 | 18.4 | 3 | 64.9 | 16.3 | 3 | 1.05 | 0.100 |
| Soil C stock (0 – 30 cm, freshwater) | 104 | 18.4 | 3 | 64.9 | 16.3 | 3 | 1.05 | 0.100 |
| CO₂-eq emissions (all wetlands) | 19.2 | 7.54 | 40 | 34.9 | 10.7 | 49 | -0.24 | 0.18 |
| CO₂-eq emissions (freshwater) | 20.2 | 8.31 | 36 | 36.5 | 11.9 | 42 | -0.24 | 0.18 |
| CO₂-eq emissions (all wetlands, absence native versus non-native) | 24.3 | 14.9 | 19 | 8.65 | 6.08 | 26 | 0.32 | 0.12 |
| CO₂-eq emissions (all wetlands, presence native versus non-native) | 27.2 | 11.3 | 21 | 38.4 | 14.5 | 33 | -0.15 | 0.038 |
| CO₂-eq emissions (all wetlands, **tropical wet** climate zone)* | 0.03 | 0.008 | 2 | 14.7 | 14.6 | 4 | -0.46 | 0.53 |
| CO₂-eq emissions (all wetlands, **tropical moist** climate zone) | 38.4 | 18.6 | 4 | 164 | 99.7 | 4 | -0.77 | 0.24 |
| CO₂-eq emissions (all wetlands, **tropical dry** climate zone) | -0.12 | 0.00 | 1 | -0.25 | 0.00 | 1 | NA | NA |
| CO₂-eq emissions (all wetlands, **warm temperate moist** climate zone)* | 29.3 | 28.5 | 2 | 31.7 | 27.7 | 5 | -0.03 | 0.50 |
| CO₂-eq emissions (all wetlands, **warm temperate dry** climate zone) | 10.1 | 2.96 | 9 | 16.6 | 4.43 | 12 | -0.48 | 0.38 |
| CO₂-eq emissions (all wetlands, **boreal dry** climate zone)* | -21.0 | 26.5 | 4 | 32.1 | 25.9 | 4 | -0.88 | 0.34 |
| CO₂-eq emissions (freshwater, **tropical wet** climate zone) | 0.03 | 0.008 | 2 | 14.7 | 14.6 | 4 | -0.46 | 0.53 |
| CO₂-eq emissions (freshwater, **tropical moist** climate zone) | 36.9 | 26.2 | 3 | 163 | 141 | 3 | -0.58 | >0.99 |
| CO₂-eq emissions (freshwater, **warm temperate moist** climate zone) | 29.3 | 28.5 | 2 | 31.7 | 27.7 | 5 | -0.03 | 0.50 |
| CO₂-eq emissions (freshwater, **warm temperate dry** climate zone) | 13.2 | 2.86 | 7 | 27.2 | 4.05 | 7 | -1.42 | 0.04 |
| CO₂-eq emissions (freshwater, **boreal dry** climate zone)* | -21.0 | 26.5 | 4 | 32.1 | 25.9 | 4 | -0.88 | 0.34 |

*All wetlands were freshwater wetlands.

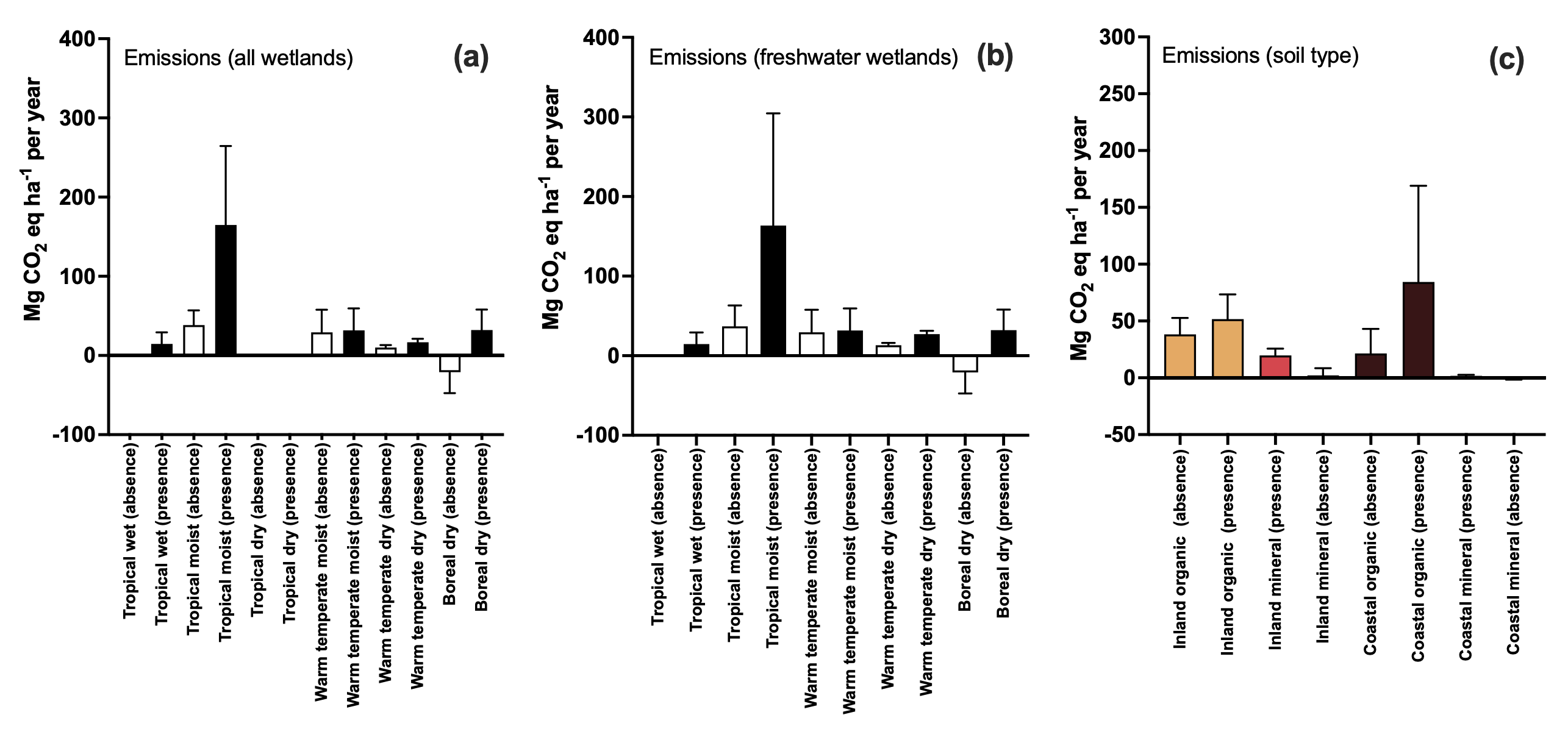

**Figure S1.** **(a)** Emissions of CO₂-eq in the absence and presence of ungulates across all wetlands and different climate zones. **(b)** Emissions of CO₂-eq in the absence and presence of ungulates across freshwater wetlands and different climate zones. **(c)** Emissions of CO₂-eq in the absence and presence of ungulates across wetlands with different soil types: inland organic, inland mineral, coastal organic, and coastal mineral soils.

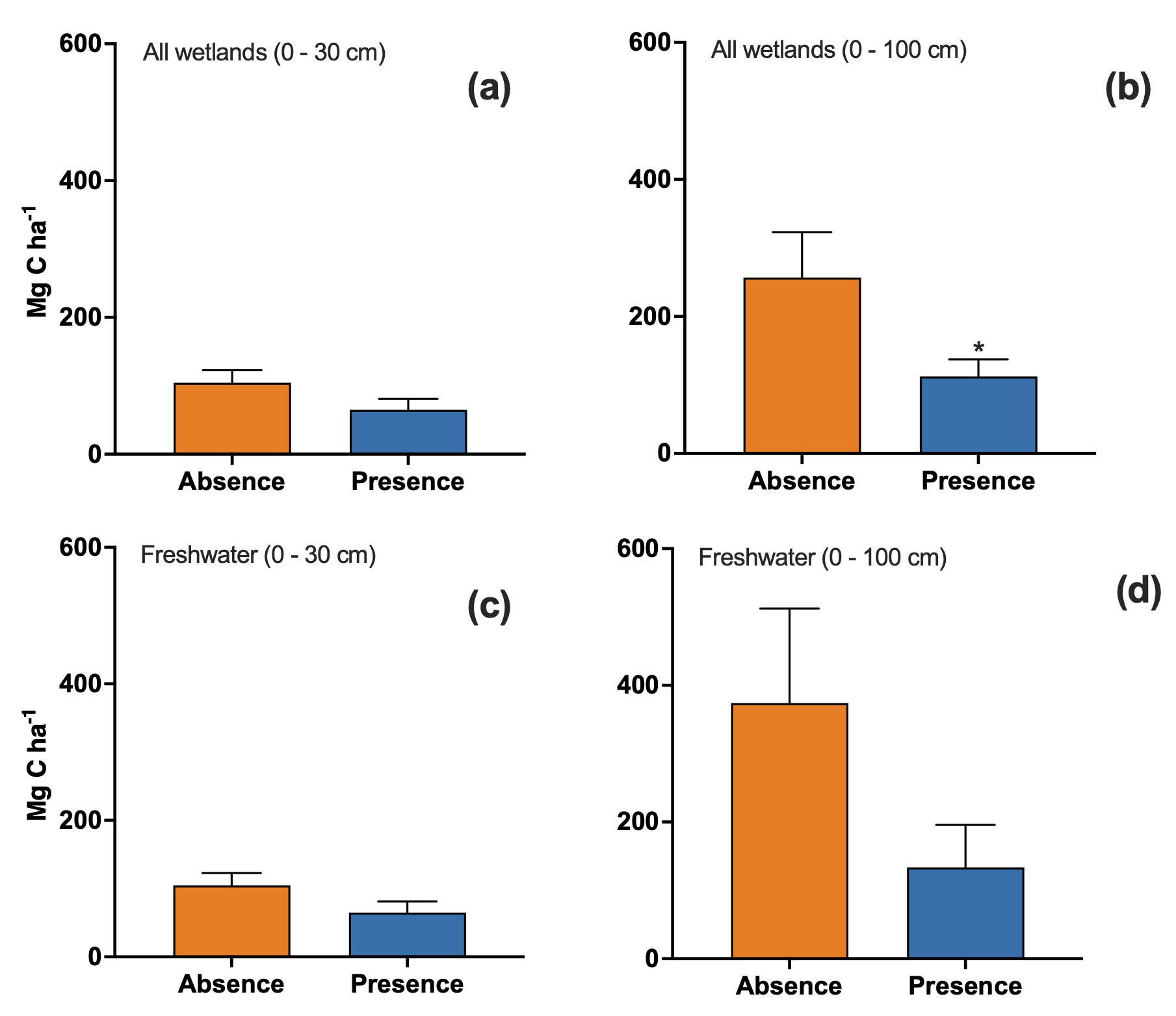

**Figure S2. (a)** Soil carbon stock (0 - 30 cm depth) in the absence and in the presence of ungulates (all wetlands were freshwater wetlands, Hedges’ *g=*1.05, *p*= 0.10). **(b)** Soil carbon stock (0 – 100 cm depth) in the absence and presence of ungulates (Hedges’*g=*0.66, *p*= 0.029). **(c)** Soil carbon stock (0- 30 cm depth) in the absence and in the presence of ungulates across freshwater wetlands (Hedges’ *g=*1.05, *p*= 0.10). **(d)** Soil carbon stock (0 - 100 cm depth) in the absence and in the presence of ungulates across freshwater wetlands (Hedges’ *g=*0.75, *p*= 0.052).
